# Mechanistic Insights into Cross-β Sheet Formation in the HIV-Associated Amyloidogenic Peptide PAP248–286 from Unbiased All-Atom Molecular Dynamics Simulation

**DOI:** 10.1101/2025.09.02.673649

**Authors:** Nikhil Agrawal, Emilio Parisini

## Abstract

Semen-derived enhancer of viral infection (SEVI) fibrils, assembled from the peptide fragment PAP248–286, enhance HIV transmission by promoting viral attachment to host cells. However, the molecular basis of SEVI nucleation and early aggregation remains unclear. Here, we conducted 80 independent all-atom molecular dynamics (MD) simulations spanning a total of 40 μs, together with 100 independent steered MD and umbrella sampling runs, to explore the dimerization and dissociation of PAP248–286 Our results show that hydrogen bonding is the dominant stabilizing force driving β-sheet formation during peptide association. Residue-level analyses identified Arg10, Val17, Glu19, and Ile20 as key contributors to inter-peptide binding, consistent with steric zipper motifs described in other amyloid systems. Steered MD revealed mechanically resilient dimers with average rupture forces of ∼20 kcal/mol/Å and multi-barrier unbinding behavior. Umbrella sampling estimated a peptide dissociation free energy of ∼8.7 kcal/mol, highlighting a clear thermodynamic separation between bound and unbound states. Together, these findings suggest that small β-sheet nuclei in PAP248–286 dimers act as cooperative intermediates that seed the formation of full-length cross-β structures, providing atomistic insights into the earliest steps of SEVI fibril assembly.

## 1. Introduction

Over the past four decades, HIV infection has emerged as a critical global health challenge. According to the World Health Organization, HIV has claimed an estimated 42.3 million lives to date, with approximately 40 million individuals living with HIV infection by the end of 2023, including 20.5 million women^1-2^. Unprotected sexual contact remains one of the primary routes of HIV transmission, with male seminal fluid serving as a key vector for viral spread to females^3-4^. A significant factor contributing to this transmission is human prostatic acid phosphatase (PAP), a protein abundant in seminal fluid, which undergoes proteolytic cleavage to generate the peptide fragment PAP248-286^5^. These peptides aggregate to form amyloid fibrils known as SEVI (Semen-derived Enhancer of Viral Infection). Due to their highly cationic nature, SEVI fibrils facilitate viral attachment to mammalian cells, and have been suggested to enhance HIV infection rates by up to 400,000-fold during sexual transmission^5-8^.

Given that amyloid formation is implicated in nearly 40 human diseases^9-10^, understanding the structural properties of SEVI fibrils is crucial. Despite sequence variations among amyloidogenic proteins, amyloid fibrils commonly adopt a cross β-sheet-rich structure^11^. The fibrillation process involves a series of structural transitions, including intermediate states such as oligomers and protofibrils^12^. Among these intermediates, structured oligomers containing cross β-sheet motifs act as nucleation centers for mature fibril formation^13-14^. Notably, in neurodegenerative disorders such as Alzheimer’s disease, oligomers are often regarded as the most toxic species^15^. Similarly, a study by Münch *et al*^5^. confirmed that SEVI amyloids adopt a cross β-sheet structure, while Usmani *et al*.^16^, using atomic force microscopy (AFM), further demonstrated that SEVI forms a dense fibrillar network in human semen, with individual fibrils measuring 0.5–2□ μm in length and 5–20□ nm in diameter— closely resembling those formed *in vitro*. Understanding how these fibrils assemble is critically important, as it provides insights into the molecular mechanisms underlying SEVI’s function and its ability to enhance viral infection. Despite these findings, no experimental structure of SEVI in its oligomeric or fibrillar state is currently available. In this context, MD simulations can play a pivotal role by modeling the self-assembly process at atomic detail, predicting transient intermediates, and offering structural insights that are difficult to capture experimentally. This integrative approach can bridge existing gaps in our understanding of SEVI’s amyloid architecture and functional behaviour.

Dimer formation is a critical step in amyloidogenesis, acting as a nucleation point for higher-order aggregation. Amyloid formation is believed to follow a pathway common to many amyloidogenic proteins, where the transition from monomers to dimers represents an early and essential stage of fibril assembly. Literature reports suggest that dimerization of amyloidogenic peptides is driven by intermolecular hydrogen bonding and hydrophobic interactions, facilitating β-sheet stabilization and the subsequent formation of higher-order oligomers^10-11^. Several studies indicate that amyloid dimers initially adopt loosely structured conformations, which gradually reorganize into stable β-sheet-rich intermediates^11-12^. These findings align with general amyloidogenesis mechanisms, wherein dimers and oligomers act as metastable species governing fibril growth kinetics^17-18^. Therefore, understanding dimerization of PAP248-286 peptides is important for uncovering structural transition pathways, which could help in developing inhibitors aimed at blocking SEVI-mediated pathological processes.

## 2. Methods

In the present study, we have performed all-atom MD simulations to understand dimer formation of PAP248-286 peptides in five different forms, based on different initial secondary structure content in PAP248-286 (Table 1).

**Table 1.**
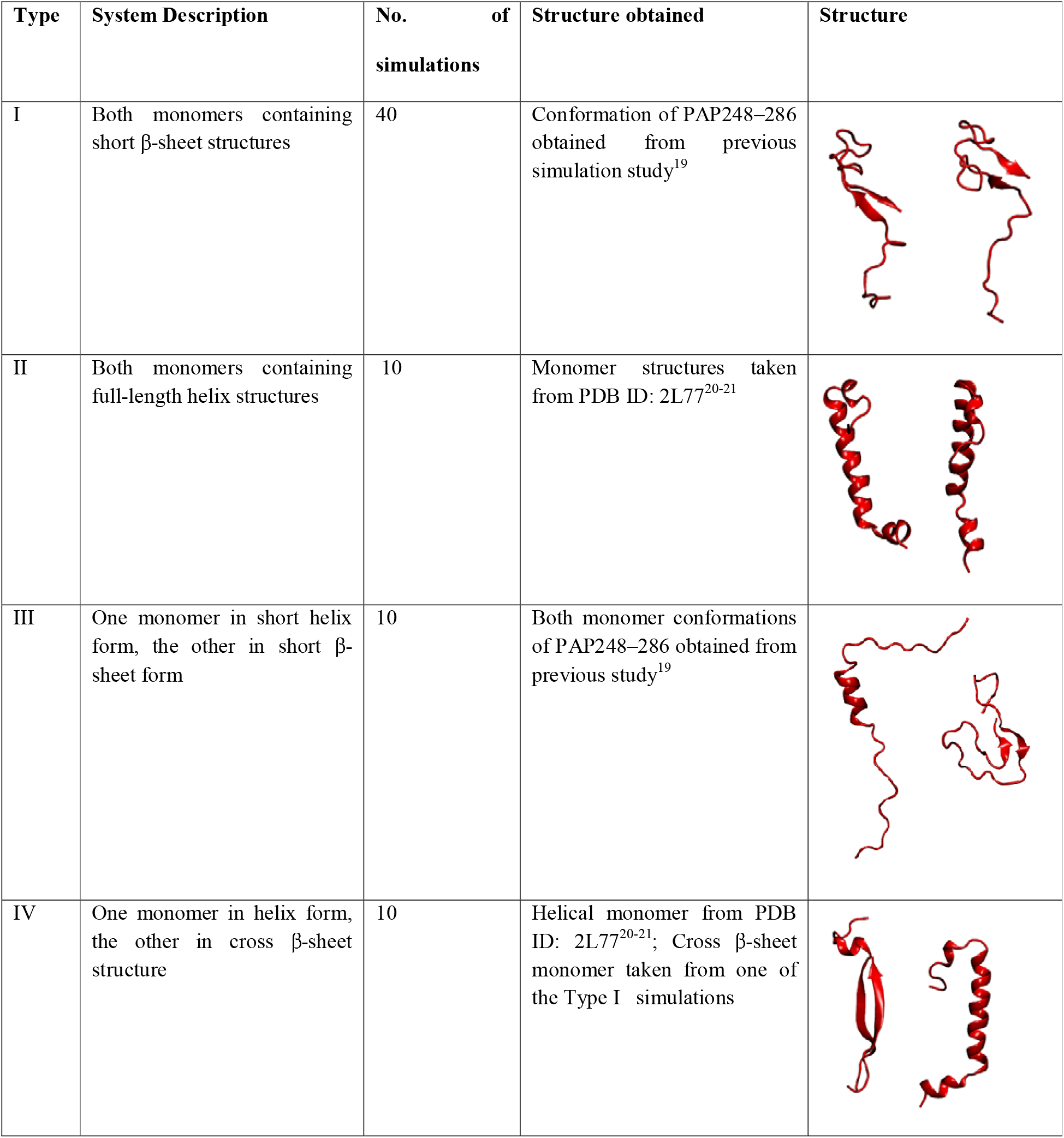
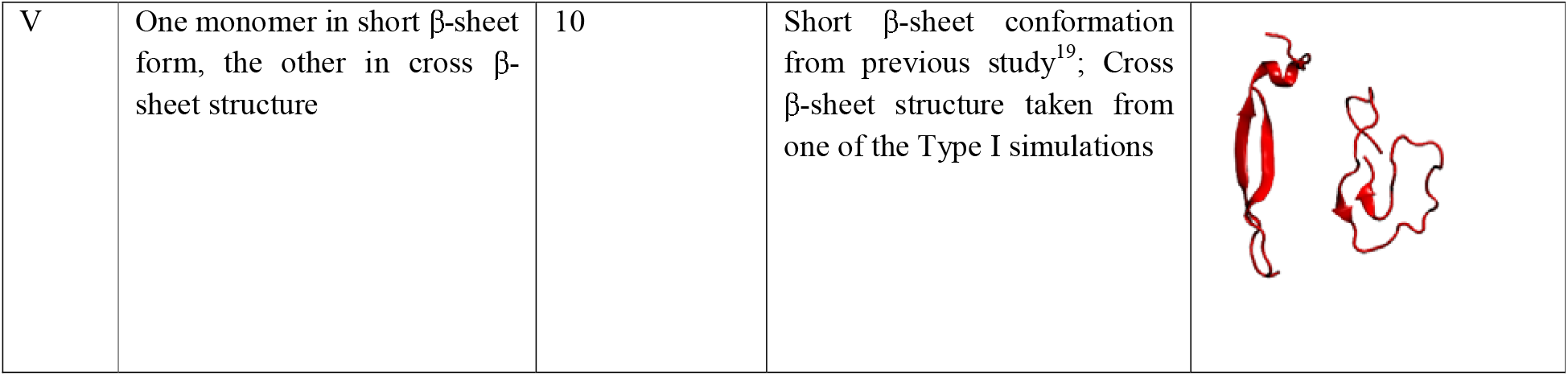
The five different simulation types performed in this study

| Type | System Description | No. of simulations | Structure obtained | Structure |
| --- | --- | --- | --- | --- |
| I | Both monomers containing short β-sheet structures | 40 | Conformation of PAP248–286 obtained from previous simulation study<sup>19</sup> | |
| II | Both monomers containing full-length helix structures | 10 | Monomer structures taken from PDB ID: 2L77<sup>20-21</sup> |  |
| III | One monomer in short helix form, the other in short β-sheet form | 10 | Both monomer conformations of PAP248–286 obtained from previous study<sup>19</sup> | |
| IV | One monomer in helix form, the other in cross β-sheet structure | 10 | Helical monomer from PDB ID: 2L77<sup>20-21</sup>; Cross β-sheet monomer taken from one of the Type I simulations | |
| V | One monomer in short β-sheet form, the other in cross β-sheet structure | 10 | Short β-sheet conformation from previous study<sup>19</sup>; Cross β-sheet structure taken from one of the Type I simulations | |

In this study, we employed the NMR structure of PAP248–286 (PDB ID: 2L77, BMRB ID: 17346) together with conformations generated in our previous simulations^19-21^. At pH 4.2, the histidine residues His3 and His23 were protonated, yielding an overall peptide charge of +8. Hydrogen atoms were subsequently added with the H++ server^22^ to reproduce the acidic vaginal environment. All molecular dynamics simulations were performed using the CHARMM36m force field^23^, an updated version of CHARMM36^24^ optimized for intrinsically disordered proteins and peptides.

Five distinct systems were constructed for simulation. Type I consisted of two PAP248–286 monomers in short β-sheet conformations solvated in 10,207 water molecules with 8 Cl□ counter ions. Type II contained two monomers in fully helical conformations solvated in 14,013 water molecules with 8 Cl□ counter ions. Type III comprised one monomer in a short helix and the other in a short β-sheet conformation, solvated in 17,489 water molecules with 8 Cl□ counter ions. Type IV contained one monomer in helical form and the other in a cross β-sheet structure, solvated in 11,265 water molecules with 8 Cl□ counter ions. Finally, Type V contained one monomer in short β-sheet and the other in cross β-sheet form, solvated in 19,641 water molecules with 8 Cl□ counter ions.

All systems were subjected to energy minimization using the steepest descent algorithm^25^, followed by two equilibration phases of 500 ps each under NVT and NPT ensembles. During minimization and the first equilibration phase, positional restraints of 400 kJ mol□ ^1^nm □ ^2^ were applied to backbone atoms and 40 kJ mol□ ^1^ nm□ ^2^ to sidechain atoms. In the second equilibration stage, restraints were applied only to backbone atoms. Production simulations were subsequently performed without restraints. Bond constraints for proteins were applied using the LINCS algorithm^26^, and water geometries were maintained with the SETTLE algorithm^27^. Long-range electrostatics were treated using the Particle Mesh Ewald (PME) method^28^, while van der Waals and short-range electrostatic interactions were truncated at 12 Å. The Nose–Hoover thermostat^29^ and Parrinello–Rahman barostat^30^ were used to maintain temperature and pressure at 310.15 K and 1 bar, respectively, each with a coupling constant of 1 ps. Integration of the equations of motion was performed with the leap-frog algorithm^31^.

For the Type I system, 40 independent production simulations were performed, while 10 simulations were conducted for each of the remaining systems. Each trajectory was 500 ns in length, yielding a total of 80 simulations and an aggregate sampling time of 40 μs. Independent replicates were initiated from randomized velocity seeds. All simulations were carried out using GROMACS^32^.

### Steered molecular dynamics (SMD) Simulations, Umbrella Sampling, and Potential of Mean Force (PMF) Calculations

To probe the mechanical stability of PAP248–286 dimers, SMD simulations were performed. A representative dimer conformation with the maximum number of inter-peptide hydrogen bonds was selected as the starting structure. One monomer was fixed, and a constant-velocity pulling force was applied to the center of mass of the second monomer along a defined reaction coordinate. A total of 100 independent SMD trajectories were generated and the maximum dissociation force (F□ □ □), dissociation time (t_d), and displacement at rupture (d) were recorded for each trajectory. Force–time and displacement–time profiles were analyzed to characterize the multi-barrier unbinding process and assess the mechanical resilience of the β-sheet interface. To quantify the thermodynamic stability of the dimer, umbrella sampling was performed along the center-of-mass (COM) distance between the two peptides. Fifty-five windows spaced by 1.0 Å covered a COM distance range of 0–55 Å. Each window was equilibrated and sampled extensively, ensuring ≥20–30% overlap between adjacent windows. Harmonic restraints maintained the COM separation in each window. The PMF was reconstructed using the weighted histogram analysis method (WHAM), and the binding free energy (ΔG_bind) was calculated as the free energy difference between the bound and unbound states, providing a quantitative measure of dimer stability.

### Analysis

Hydrogen bonds and minimum inter-peptide distances were calculated using the GROMACS^32^ tools gmx hbond and gmx mindistance, respectively. Binding free energies and per-residue contributions were evaluated using the gmx_MMPBSA tool^33^. In the SMD simulations, force–distance profiles were analyzed to identify rupture events and characterize the strength and dynamics of peptide–peptide interactions. For umbrella sampling simulations, the WHAM was used to compute the PMF profiles along the chosen reaction coordinates.

## 3. Results

### 3.1 Interaction between two PAP248-286 monomers and change in conformation

Hydrogen bonding is a key determinant of protein–protein interactions, particularly in amyloid-forming systems, where it stabilizes extended β-sheet structures and facilitates the assembly of ordered aggregates^34^. Previous studies have indicated that inter-peptide distances below ∼6 Å are required to enable backbone–backbone hydrogen bonding^35^, which is critical for maintaining β-sheet conformations and forming the cross-β spines characteristic of amyloid fibrils. In our study, across 40 independent simulations of the Type-I set, we observed multiple trajectories showing an increase in β-sheet content in both peptides upon interaction. Notably, this effect was unique to the Type-I simulations, as the other sets did not exhibit full cross β-sheet formation during peptide association (Supplementary Fig. 3–6). Consequently, our analysis focuses primarily on the Type-I data.

Fig. 1 presents the time evolution of hydrogen bonds, β-sheet content; inter monomer distance, and binding free energy. In a representative trajectory, the minimum distance between the two monomers dropped below 5 Å around 400 ns (Fig. 1C), this distance, coinciding with a sharp increase in the number of hydrogen bonds formed between the peptides (Fig. 1A). This event was accompanied by a simultaneous rise in the number of residues adopting β-sheet conformations (Fig. 1B), reflecting enhanced structural ordering. Consistently, binding energy calculations (Fig. 1D) indicated a highly favorable interaction during the 400–450 ns interval, suggesting the establishment of a stable binding event. Fig. 2 illustrates the evolution of secondary structure content in the PAP248–286 monomers at different time points, highlighting the structural rearrangements induced by peptide association.

**Fig. 1.**
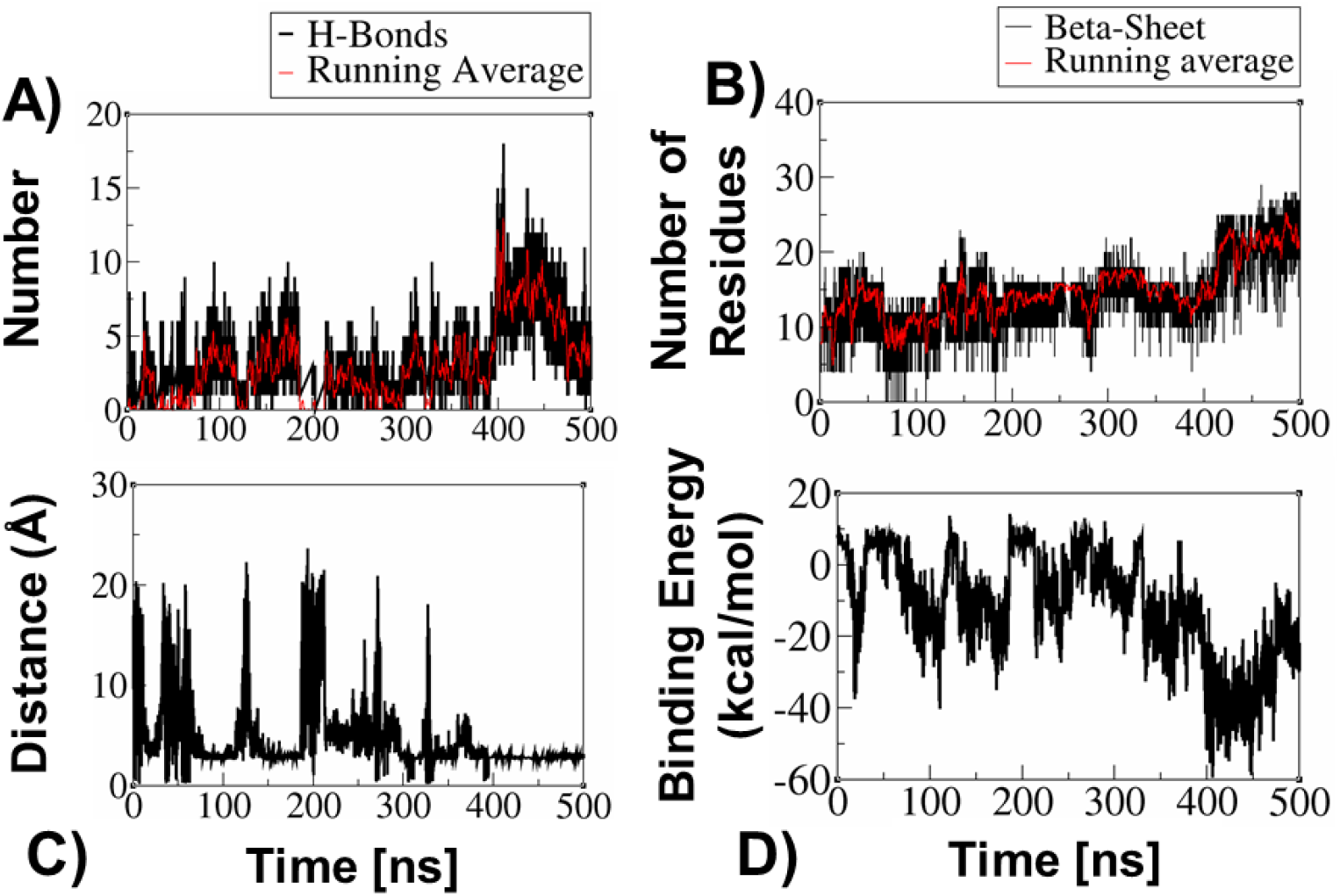
A) Time evolution of the number of H-Bonds, B) Time evolution of the number of residues of PAP248-826 members in β-sheet conformation, C) Minimum distance between two PAP248-286 monomers, and D) Time evolution of the binding energy between two PAP248-286 monomers.

**Fig. 2.**
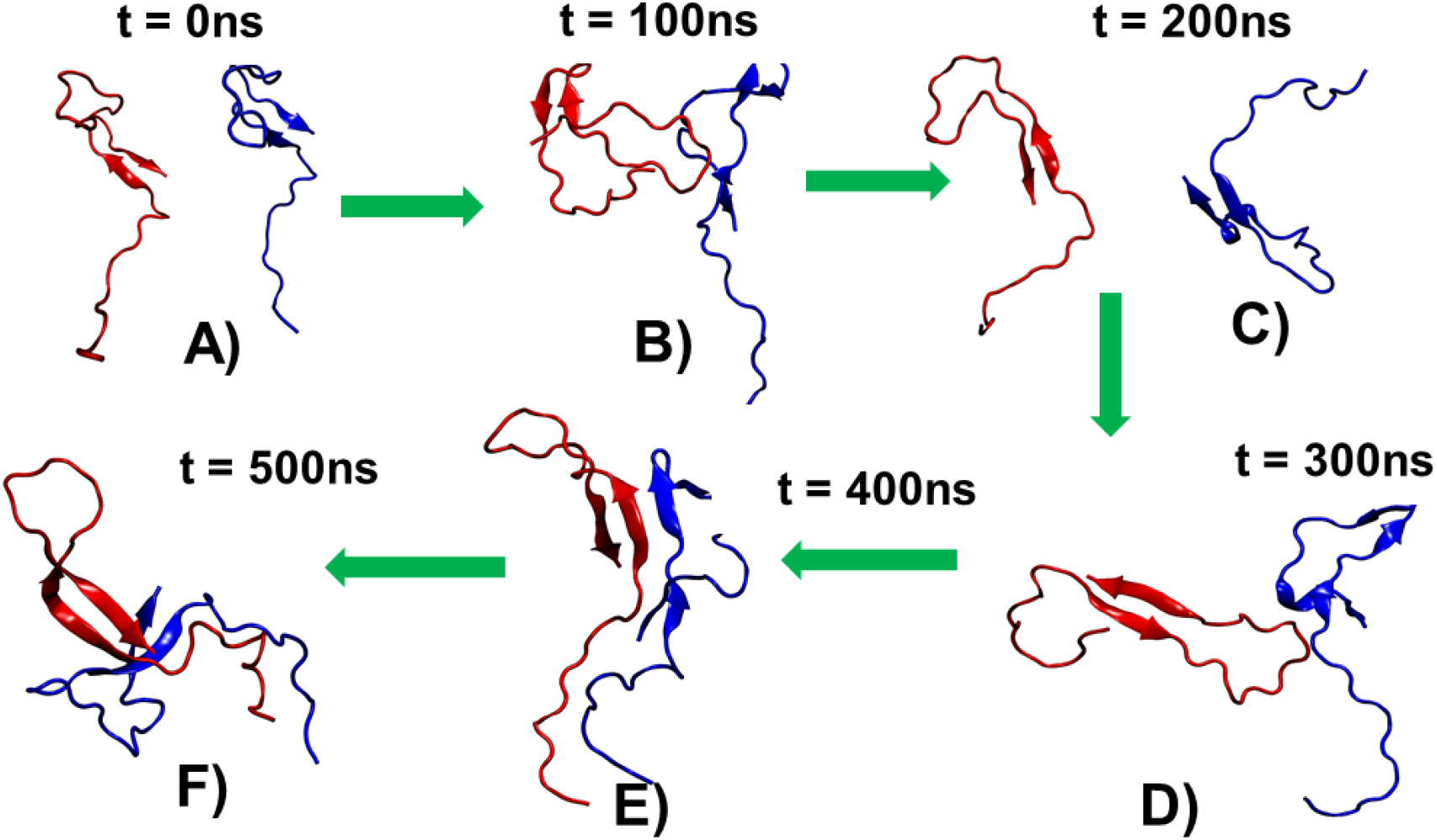
PAP248-286 monomer structures from one of the representative trajectory at six different time points, A) t = 0 ns, B) t = 100 ns, C) t = 200 ns, D) t = 300 ns, E) t = 400 ns, and F) t = 500 ns.

Overall, these findings suggest that the close approach of peptide monomers facilitates the formation of hydrogen bonds, which play a pivotal role in stabilizing β-sheet structures. This stabilization promotes the nucleation and growth of amyloid assemblies by driving the formation of stable, ordered aggregates, providing molecular-level insight into the early stages of amyloid fibril formation.

### 3.2 Identification of residues involved in the binding of PAP248-286 monomers

Several studies have shown that specific interactions between residues of one monomer and those of another play a central role in amyloid peptide assembly and promote β-sheet formation and stacking^9,36^. In bacterial amyloids such as CsgA and TasA, conserved segments like LNIYQY and VTQVGF mediate cross-β interactions, where residues from one peptide align precisely with those from another, forming tightly interdigitated steric zippers^37-38^.

To identify the residues and regions of each monomer involved in binding, we calculated the minimum inter-peptide distances during the 400–425 ns window—when binding energies were most favorable (Fig. 3D). Additionally, we computed the time evolution of the per-residue binding energy contributions during this period. The per-residue distance plot (Fig. 3A) revealed that residues 6 to 24 from both peptides were in closest proximity during this time.

**Fig. 3.**
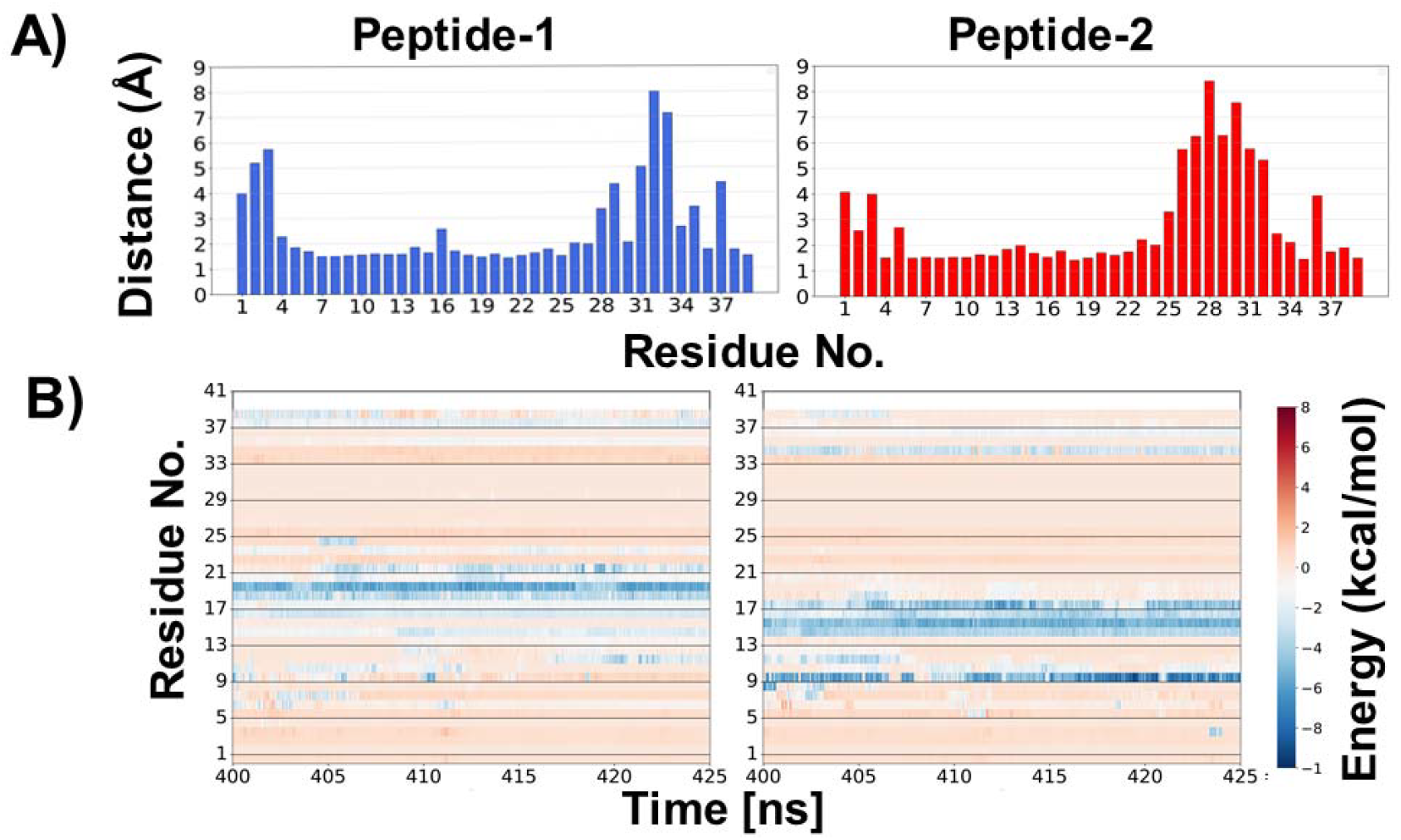
A) Minimum distance between both peptide during 400-425 ns, B) Time evolutions of contribution of each residue in binding.

To further pinpoint key contributors to binding, we analyzed the per-residue binding energy contributions between 400 and 425 ns (Fig. 3B). In the first peptide, residues 10 to 21 showed the highest contributions, while in the second peptide, residues 10 to 18 were most energetically involved. Among these, Glu19 and Ile20 from the first peptide, and Arg10, Gly14, Val15, Leu16, Val17, and Asn18 from the second peptide, played prominent roles in the interaction. These residues engage in both hydrophobic and electrostatic interactions, highlighting the importance of both types of forces in stabilizing the peptide-peptide interface.

### 3.3 SMD Simulations force and peptide dissociation

To investigate the strength of association between two monomers, we examined the unbinding mechanisms of two PAP248-826 monomers. We conducted 100 independent SMD simulations, applying constant-velocity pulling to dissociate spontaneously associated peptide dimers (Fig. 4). For the SMD simulations, we selected a starting conformation of the PAP248-286 dimer that contained the most inter peptide hydrogen bonds. From SMD simulations, we extracted three key descriptors: the maximum dissociation force (F□ □ □), the dissociation time (t_d), and the displacement at unbinding (d). The force–time profiles (Fig. 4A) exhibited a characteristic pattern in which the pulling force gradually increased, peaking at an average F□ □ □ of 20.29 ± 1.90 kcal/mol/Å, before abruptly dropping at the dissociation event. This magnitude of force reflects the mechanical stability of the peptide complex, underscoring the strength of β-sheet-mediated inter-peptide interactions. Such mechanical robustness is consistent with previous AFM studies on amyloid fibrils, which have shown that β-sheet-rich aggregates resist dissociation forces in the nanonewton (nN) range^39-40^, and highlights the biological relevance of these structures in maintaining amyloid integrity under stress. The mean dissociation time (t_d) was found to be 124.73 ± 20.07 ps, with noticeable variability across trajectories, suggesting a rugged energy landscape with multiple intermediate states or alternative dissociation pathways. This observation aligns with earlier computational findings that amyloid peptide dissociation does not follow a single-path mechanism, but rather involves the crossing of multiple kinetic barriers. Furthermore, the displacement-time profiles (Fig. 4B) revealed an initial elastic extension of the complex followed by a sharp increase in displacement, marking the rupture point. The average displacement at the moment of dissociation was 8.10 ± 1.91 Å, corresponding to the disruption of backbone hydrogen bonds and side-chain contacts that stabilize the β-sheet interface.

**Fig. 4.**
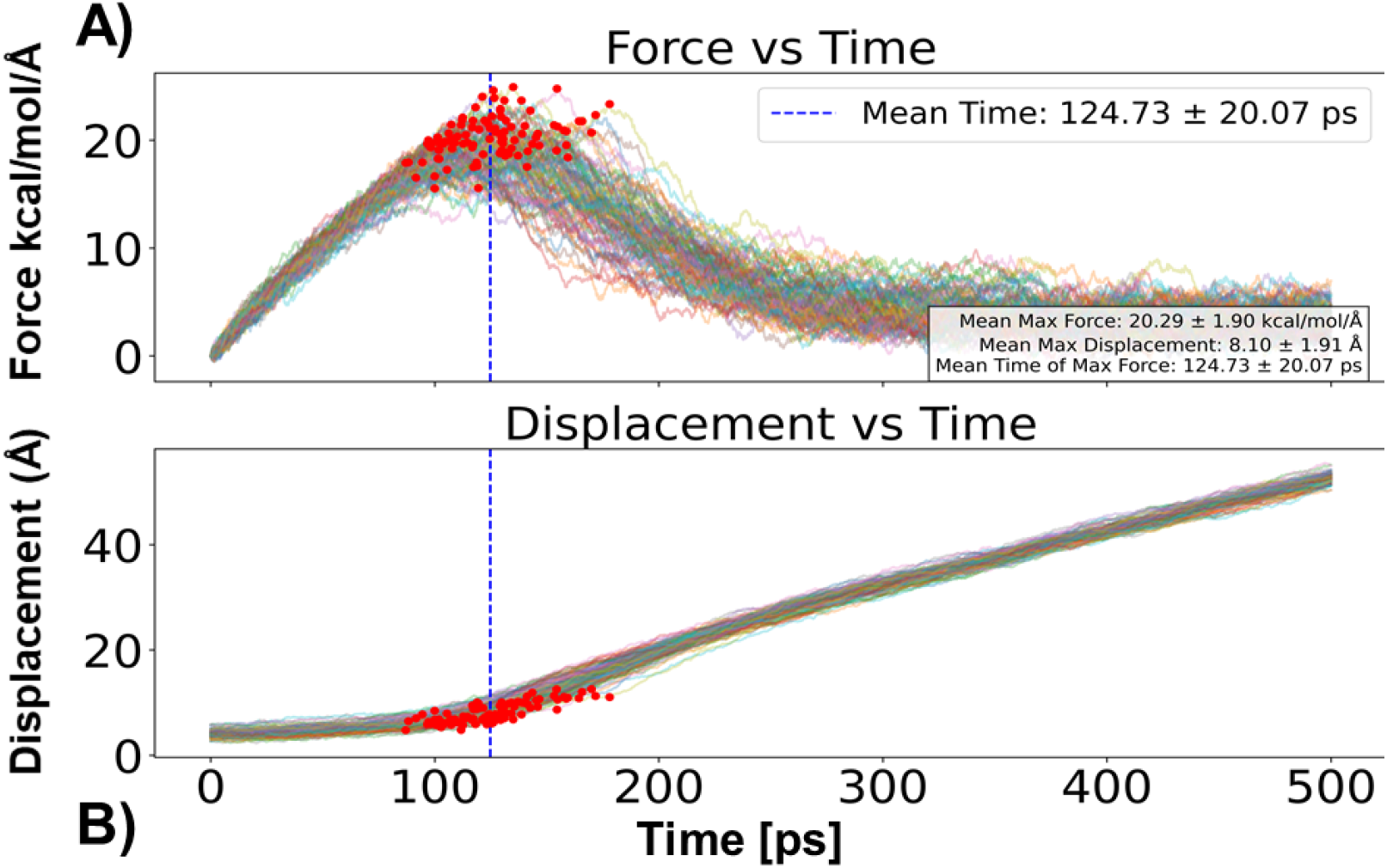
A) force vs time graph for all 100 SMD simulations, B) displacement vs time for 100 SMD simulations.

Importantly, we observed a strong positive correlation (R^2^ > 0.85) between F□ □ □ and t_d, indicating that complexes requiring higher dissociation forces also took longer to separate. This suggests a force-dependent, multi-step unbinding mechanism in which higher-energy barriers delay dissociation. Such behavior reinforces the notion that amyloid dissociation is governed more by kinetic control than by a single thermodynamic transition state. This is particularly significant given the role of β-sheets conformation in various physiological and pathological contexts, such as biofilm formation in bacteria or the persistence of neurotoxic aggregates in neurodegenerative diseases. The mechanical resistance of these assemblies, as captured in our SMD simulations, provides molecular insight into the interaction strength between two monomers of PAP248-286 in β-sheet conformation.

Taken together, these results highlight the mechanically robust and kinetically complex nature of PAP248-286 peptides dissociation. The combination of force-resistance, displacement behavior, and variable dissociation timing underscores the cooperative nature of PAP248-286 monomer unbinding in β-sheet form as peptide dissociated β-sheet structure was reduced as appeared to be in conformation similar as it was at the beginning of the MD simulation (Fig, 5). By providing atomistic-level understanding of dissociation pathways, our study contributes to the broader understanding of amyloid stability, with potential implications for designing molecules that can modulate or disrupt pathogenic peptide aggregation.

**Fig. 5.**
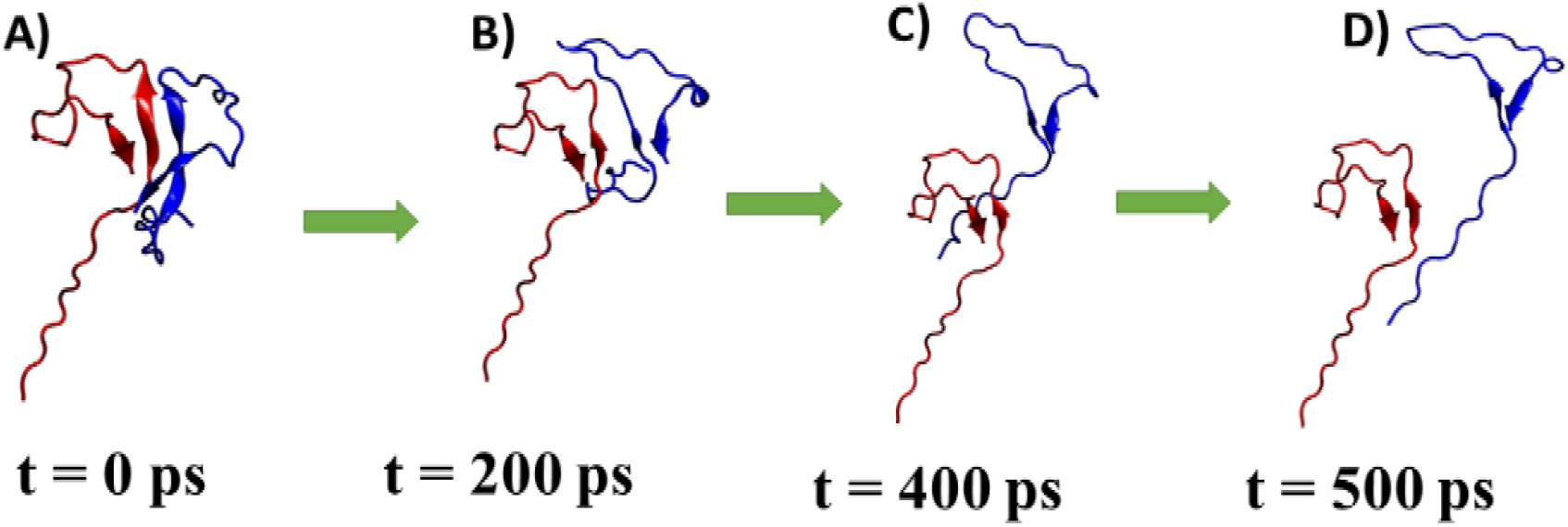
A) Snapshots of PAP248-286 monomers dissociation from one of the SMD trajectories, A) t = 0 ps, B) t = 200 ps, C) t = 400 ps, D) t = 500 ps.

### 3.4 Umbrella Sampling and PMF Analysis of Peptide Dissociation

To quantify the free energy profile associated with the dissociation of the peptide dimer, we performed umbrella-sampling simulations using a series of harmonic restraints along the center-of-mass (COM) distance between the two peptides. A total of 55 windows were generated with a spacing of 1.0 Å, covering a COM distance range from 0 to 55 Å. Each window was equilibrated and sampled extensively to ensure adequate overlap of the restrained configurations.

Fig. 6A illustrates the probability distribution of the reaction coordinate sampled in each umbrella window. The histograms demonstrate sufficient overlap between adjacent windows (≥ 20–30%), indicating robust sampling across the entire reaction coordinate. This overlap is critical for reliable reconstruction of PMF via WHAM.

**Fig. 6.**
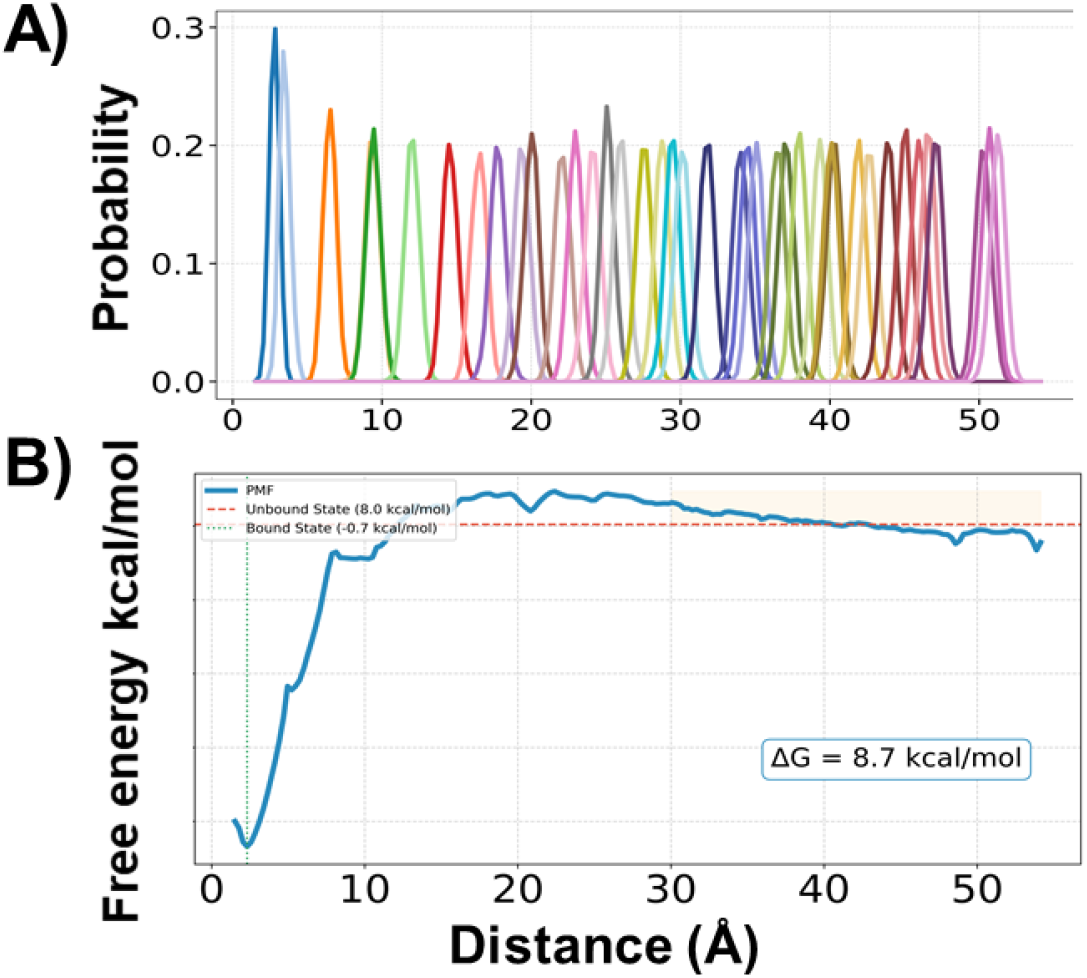
A) Probability distributions from each umbrella window, B) PMF profile indicating a bound state and unbound state of PAP248-286 monomers

The resulting PMF curve is shown in Fig. 6B. A distinct global minimum is observed at short COM distances, corresponding to the bound state of the peptide dimer. The free energy gradually increases with separation, plateauing at larger distances that correspond to the unbound state. The bound state is located at approximately 3 Å with a free energy of -0.7 kcal/mol, while the unbound state stabilizes at ∼8.0 kcal/mol. The calculated binding free energy (ΔG_bind) is 8.7 kcal/mol, suggesting a moderately strong interaction between the two peptides in the dimeric state. This PMF profile provides critical insight into the thermodynamic stability of the dimer and supports the hypothesis that dimer formation is a key intermediate step during peptide aggregation.

## 4. Discussion and Conclusion

Our study provides an integrated view of the molecular determinants and energetics underlying the dimerization and dissociation of PAP248-286 peptides, a process central to SEVI fibril formation and its role in enhancing HIV infectivity. Through a combination of unbiased MD simulations residue-level interaction analysis, SMD, and umbrella sampling, we reveal how inter-peptide interactions drive conformational transitions critical to amyloid nucleation. Hydrogen bonding between both monomers played a dominant stabilizing force, facilitating β-sheet formation upon peptide association. This aligns with previous structural models of amyloid fibrils, where inter-strand hydrogen bonds and side-chain contacts form highly ordered, β-sheet-rich assemblies^34^. Notably, the residues identified as key contributors to binding—such as Glu19, Ile20, Arg10, and Val17—mirror interaction motifs observed in other amyloid systems like CsgA, which rely on steric zipper arrangements for stability^41-42^. The mechanical stability of the dimer, as revealed by SMD simulations, is comparable to that observed in AFM studies of amyloids, with force–time and displacement–time profiles indicating a multi-barrier unbinding landscape^43-44^. The observed strong correlation between unbinding force and dissociation time supports a kinetic model of dissociation, where rupture is governed not by a single transition state but by cooperative disruption of backbone and side-chain interactions—consistent with findings in other amyloid-forming system. Thermodynamic analysis via umbrella sampling corroborates these findings. The calculated dissociation free energy (∼8.7 kcal/mol) and clear separation between bound and unbound states confirm that dimer formation is a moderately favorable and distinct intermediate along the aggregation pathway. The significant histogram overlap further validates the robustness of the free energy estimation. Collectively, these observations support a model in which PAP248-286 dimerization involves structurally and energetically cooperative interactions that stabilize β-sheet conformations. This structural transition, triggered by proximity and residue-specific contacts, likely serves as a nucleating event for higher-order aggregation. Given SEVI’s enhancement of HIV infection, these insights offer a molecular rationale for developing small molecules or peptides to disrupt early dimer formation and potentially modulate amyloid-related pathologies.

## Acknowledgements

N.A. acknowledges the RRF grant No. 65/OSI/PG (RRF project No. 5.2.1.1.i.0/2/24/I/CFLA/001) for funding, and E.P. thanks the Latvian Recovery and Resilience Fund (grant No. 74/OSI/ZG) for financial support. We are grateful to the Latvian Institute of Organic Synthesis, Riga, for providing supercomputing resources. N.A. further acknowledges the Centre for High Performance Computing (CHPC), Cape Town, for access to computational facilities.

## References

1. Multani, A.; Becken III, B.; Padival, S.; Cunningham, C. K., Human Immunodeficiency Virus I: History, Epidemiology, Transmission, and Pathogenesis. In Introduction to Clinical Infectious Diseases: A Problem-Based Approach, Springer: 2025; pp 661–667.

2. Tang, L.; Du, Y.-T.; Kong, W.-H.; Liu, P.; Zhu, Z.-R.; Xie, S.-Z.; Liu, M.-Q., Late HIV/AIDS diagnosis among people living with HIV in Wuhan in 2023. Frontiers in Microbiology 2025, 16, 1594847.

3. Nambiar, P.; Short, W. R., Mechanisms of HIV transmission. Fundamentals of HIV Medicine 2019, 8, 20.

4. Sabatté, J.; Lenicov, F. R.; Cabrini, M.; Rodrigues, C. R.; Ostrowski, M.; Ceballos, A.; Amigorena, S.; Geffner, J., The role of semen in sexual transmission of HIV: beyond a carrier for virus particles. Microbes and infection 2011, 13 (12-13), 977–982.

5. Münch, J.; Rücker, E.; Ständker, L.; Adermann, K.; Goffinet, C.; Schindler, M.; Wildum, S.; Chinnadurai, R.; Rajan, D.; Specht, A., Semen-derived amyloid fibrils drastically enhance HIV infection. Cell 2007, 131 (6), 1059–1071.

6. Lee, Y. H.; Ramamoorthy, A., Semen-derived amyloidogenic peptides—Key players of HIV infection. Protein science 2018, 27 (7), 1151–1165.

7. Kim, K.-A.; Yolamanova, M.; Zirafi, O.; Roan, N. R.; Staendker, L.; Forssmann, W.-G.; Burgener, A.; Dejucq-Rainsford, N.; Hahn, B. H.; Shaw, G. M., Semen-mediated enhancement of HIV infection is donor-dependent and correlates with the levels of SEVI. Retrovirology 2010, 7, 1–12.

8. Elias, A. K.; Scanlon, D.; Musgrave, I. F.; Carver, J. A., SEVI, the semen enhancer of HIV infection along with fragments from its central region, form amyloid fibrils that are toxic to neuronal cells. Biochimica et Biophysica Acta (BBA)-Proteins and Proteomics 2014, 1844 (9), 1591–1598.

9. Chiti, F.; Dobson, C. M., Protein misfolding, functional amyloid, and human disease. Annu. Rev. Biochem. 2006, 75 (1), 333–366.

10. Chiti, F.; Dobson, C. M., Protein misfolding, amyloid formation, and human disease: a summary of progress over the last decade. Annual review of biochemistry 2017, 86 (1), 27–68.

11. Knowles, T. P.; Vendruscolo, M.; Dobson, C. M., The amyloid state and its association with protein misfolding diseases. Nature reviews Molecular cell biology 2014, 15 (6), 384–396.

12. Cohen, S. I.; Linse, S.; Luheshi, L. M.; Hellstrand, E.; White, D. A.; Rajah, L.; Otzen, D. E.; Vendruscolo, M.; Dobson, C. M.; Knowles, T. P., Proliferation of amyloid-β42 aggregates occurs through a secondary nucleation mechanism. Proceedings of the National Academy of Sciences 2013, 110 (24), 9758–9763.

13. Bieschke, J.; Herbst, M.; Wiglenda, T.; Friedrich, R. P.; Boeddrich, A.; Schiele, F.; Kleckers, D.; Lopez del Amo, J.M.; Grüning, B. A.; Wang, Q., Small-molecule conversion of toxic oligomers to nontoxic β-sheet–rich amyloid fibrils. Nature chemical biology 2012, 8 (1), 93–101.

14. Liu, H.; Qian, C.; Yang, T.; Wang, Y.; Luo, J.; Zhang, C.; Wang, X.; Wang, X.; Guo, Z., Small molecule-mediated co-assembly of amyloid-β oligomers reduces neurotoxicity through promoting non-fibrillar aggregation. Chemical Science 2020, 11 (27), 7158–7169.

15. Rinauro, D. J.; Chiti, F.; Vendruscolo, M.; Limbocker, R., Misfolded protein oligomers: Mechanisms of formation, cytotoxic effects, and pharmacological approaches against protein misfolding diseases. Molecular Neurodegeneration 2024, 19 (1), 20.

16. Usmani, S. M.; Zirafi, O.; Müller, J. A.; Sandi-Monroy, N. L.; Yadav, J. K.; Meier, C.; Weil, T.; Roan, N. R.; Greene, W. C.; Walther, P., Direct visualization of HIV-enhancing endogenous amyloid fibrils in human semen. Nature communications 2014, 5 (1), 3508.

17. Eisenberg, D.; Jucker, M., The amyloid state of proteins in human diseases. Cell 2012, 148 (6), 1188–1203.

18. Agrawal, N.; Skelton, A. A., Structure and Function of Alzheimer’s Amyloid βeta Proteins from Monomer to Fibrils: A Mini Review. The Protein Journal 2019, 38 (4), 425–434.

19. Agrawal, N.; Parisini, E., Early stages of misfolding of PAP248-286 at two different pH values: An insight from molecular dynamics simulations. Computational and Structural Biotechnology Journal 2022, 20, 4892–4901.

20. Brender, J. R.; Nanga, R. P. R.; Popovych, N.; Soong, R.; Macdonald, P. M.; Ramamoorthy, A., The amyloidogenic SEVI precursor, PAP248-286, is highly unfolded in solution despite an underlying helical tendency. Biochimica et Biophysica Acta (BBA)-Biomembranes 2011, 1808 (4), 1161–1169.

21. Nanga, R. P.; Brender, J. R.; Vivekanandan, S.; Popovych, N.; Ramamoorthy, A., NMR structure in a membrane environment reveals putative amyloidogenic regions of the SEVI precursor peptide PAP248− 286. Journal of the American Chemical Society 2009, 131 (49), 17972–17979.

22. Gordon, J. C.; Myers, J. B.; Folta, T.; Shoja, V.; Heath, L. S.; Onufriev, A., H++: a server for estimating p K as and adding missing hydrogens to macromolecules. Nucleic acids research 2005, 33 (Suppl_2), W368–W371.

23. Huang, J.; Rauscher, S.; Nawrocki, G.; Ran, T.; Feig, M.; De Groot, B. L.; Grubmüller, H.; MacKerell Jr, A. D., CHARMM36m: an improved force field for folded and intrinsically disordered proteins. Nature methods 2017, 14 (1), 71–73.

24. Huang, J.; MacKerell Jr, A. D., CHARMM36 all-atom additive protein force field: Validation based on comparison to NMR data. Journal of computational chemistry 2013, 34 (25), 2135–2145.

25. Bixon, M.; Lifson, S., Potential functions and conformations in cycloalkanes. Tetrahedron 1967, 23 (2), 769–784.

26. Hess, B.; Bekker, H.; Berendsen, H. J.; Fraaije, J. G., LINCS: A linear constraint solver for molecular simulations. Journal of computational chemistry 1997, 18 (12), 1463–1472.

27. Miyamoto, S.; Kollman, P. A., Settle: An analytical version of the SHAKE and RATTLE algorithm for rigid water models. Journal of computational chemistry 1992, 13 (8), 952–962.

28. Darden, T.; York, D.; Pedersen, L., Particle mesh Ewald: An N log (N) method for Ewald sums in large systems. Journal of chemical physics 1993, 98, 10089–10089.

29. NOSÉ, S. I., A molecular dynamics method for simulations in the canonical ensemble. Molecular physics 2002, 100 (1), 191–198.

30. Parrinello, M.; Rahman, A., Polymorphic transitions in single crystals: A new molecular dynamics method. Journal of Applied physics 1981, 52 (12), 7182–7190.

31. Van Gunsteren, W. F.; Berendsen, H. J., A leap-frog algorithm for stochastic dynamics. Molecular Simulation 1988, 1 (3), 173–185.

32. Abraham, M. J.; Murtola, T.; Schulz, R.; Páll, S.; Smith, J. C.; Hess, B.; Lindahl, E., GROMACS: High performance molecular simulations through multi-level parallelism from laptops to supercomputers. SoftwareX 2015, 1, 19–25.

33. Valdés-Tresanco, M. S.; Valdés-Tresanco, M. E.; Valiente, P. A.; Moreno, E., gmx_MMPBSA: a new tool to perform end-state free energy calculations with GROMACS. Journal of chemical theory and computation 2021, 17 (10), 6281–6291.

34. Tsemekhman, K.; Goldschmidt, L.; Eisenberg, D.; Baker, D., Cooperative hydrogen bonding in amyloid formation. Protein science 2007, 16 (4), 761–764.

35. Jaroniec, C. P.; MacPhee, C. E.; Bajaj, V. S.; McMahon, M. T.; Dobson, C. M.; Griffin, R. G., High-resolution molecular structure of a peptide in an amyloid fibril determined by magic angle spinning NMR spectroscopy. Proceedings of the National Academy of Sciences 2004, 101 (3), 711–716.

36. Sawaya, M. R.; Sambashivan, S.; Nelson, R.; Ivanova, M. I.; Sievers, S. A.; Apostol, M. I.; Thompson, M. J.; Balbirnie, M.; Wiltzius, J. J.; McFarlane, H. T., Atomic structures of amyloid cross-β spines reveal varied steric zippers. Nature 2007, 447 (7143), 453–457.

37. Wang, X.; Smith, D. R.; Jones, J. W.; Chapman, M. R., In vitro polymerization of a functional Escherichia coli amyloid protein. Journal of Biological Chemistry 2007, 282 (6), 3713–3719.

38. Romero, D.; Aguilar, C.; Losick, R.; Kolter, R., Amyloid fibers provide structural integrity to Bacillus subtilis biofilms. Proceedings of the National Academy of Sciences 2010, 107 (5), 2230–2234.

39. Knowles, T. P.; Fitzpatrick, A. W.; Meehan, S.; Mott, H. R.; Vendruscolo, M.; Dobson, C. M.; Welland, M. E., Role of intermolecular forces in defining material properties of protein nanofibrils. science 2007, 318 (5858), 1900–1903.

40. Smith, J. F.; Knowles, T. P.; Dobson, C. M.; MacPhee, C. E.; Welland, M. E., Characterization of the nanoscale properties of individual amyloid fibrils. Proceedings of the National Academy of Sciences 2006, 103 (43), 15806–15811.

41. Perov, S.; Lidor, O.; Salinas, N.; Golan, N.; Tayeb-Fligelman, E.; Deshmukh, M.; Willbold, D.; Landau, M., Structural insights into curli CsgA cross-β fibril architecture inspire repurposing of anti-amyloid compounds as anti-biofilm agents. PLoS pathogens 2019, 15 (8), e1007978.

42. Bu, F.; Dee, D. R.; Liu, B., Structural insight into Escherichia coli CsgA amyloid fibril assembly. MBio 2024, 15 (4), e00419–24.

43. Mesquida, P.; Riener, C. K.; MacPhee, C. E.; McKendry, R. A., Morphology and mechanical stability of amyloid-like peptide fibrils. Journal of Materials Science: Materials in Medicine 2007, 18 (7), 1325–1331.

44. Watanabe-Nakayama, T.; Sahoo, B. R.; Ramamoorthy, A.; Ono, K., High-speed atomic force microscopy reveals the structural dynamics of the amyloid-β and amylin aggregation pathways. International journal of molecular sciences 2020, 21 (12), 4287.

